# Novel Insight into the Aetiology of Autism Spectrum Disorder Gained by Integrating Expression Data with Genome-wide Association Statistics

**DOI:** 10.1101/480624

**Authors:** Oliver Pain, Andrew J. Pocklington, Peter A. Holmans, Nicholas J. Bray, Heath E. O’Brian, Lynsey S. Hall, Antonio F. Pardiñas, Michael C. O’Donovan, Michael J. Owen, Richard Anney

**Author notes:** Corresponding author: Dr Richard Anney, MRC Centre for Neuropsychiatric Genetics and Genomics, Cardiff University, Hadyn Ellis Building, Cardiff, CF24 4HQ.

## Abstract

**Background:** A recent genome-wide association study (GWAS) of autism spectrum disorders (ASD) (N_cases_=18,381, N_controls_=27,969) has provided novel opportunities for investigating the aetiology of ASD. Here, we integrate the ASD GWAS summary statistics with summary-level gene expression data to infer differential gene expression in ASD, an approach called transcriptome-wide association study (TWAS).

**Methods:** Using FUSION software, ASD GWAS summary statistics were integrated with predictors of gene expression from 16 human datasets, including adult and fetal brain. A novel adaptation of established statistical methods was then used to test for enrichment within candidate pathways, specific tissues, and at different stages of brain development. The proportion of ASD heritability explained by predicted expression of genes in the TWAS was estimated using stratified linkage disequilibrium-score regression.

**Results:** This study identified 14 genes as significantly differentially expressed in ASD, 13 of which were outside of known genome-wide significant loci (±500kb). *XRN2*, a gene proximal to an ASD GWAS locus, was inferred to be significantly upregulated in ASD, providing insight into functional consequence of this associated locus. One novel transcriptome-wide significant association from this study is the downregulation of *PDIA6*, which showed minimal evidence of association in the GWAS, and in gene-based analysis using MAGMA. Predicted gene expression in this study accounted for 13.0% of the total ASD SNP-heritability.

**Conclusion:** This study has implicated several genes as significantly up-/down-regulated in ASD providing novel and useful information for subsequent functional studies. This study also explores the utility of TWAS-based enrichment analysis and compares TWAS results with a functionally agnostic approach.

## Introduction

Autism spectrum disorders (ASD) are a group of neurodevelopmental disorders characterised by impaired social and communication skills, and stereotyped and repetitive behaviours. ASD has a prevalence of 1% (1), with symptoms typically starting in early childhood. Twin studies estimate the heritability of ASD at between ∼65-90% (2,3), demonstrating that genetic differences play an important role in the development of ASD. Common genetic variants conferring individually weak effects are an important component of ASD liability, with the most recent SNP-heritability estimate for ASD being 11.8% on a liability scale (assuming prevalence of 1.2%) (4).

Genome-wide association studies (GWAS) are a powerful approach for understanding the role of common alleles in the genetic aetiology of traits and disorders, and have provided several insights into the aetiology of ASD. The most recent and largest ASD GWAS including 18,381 ASD cases and 27,969 controls reported three independent loci achieving genome-wide significance (4). Genes associated with ASD were highlighted through proximity to genome-wide significant loci and via joint statistical analysis of variants within gene regions. However, a variant’s proximity to a gene is only one metric for illuminating its functional consequence and the nearest gene often does not drive the association (5).

An additional approach for highlighting genes that underlie a GWAS association is through integration of functional data. For example, Grove *et al*. used chromatin conformation data to infer whether significant ASD-loci physically interact with the surrounding genes. Alternatively, prior knowledge of variants effecting gene expression, known as expression quantitative trait loci (eQTLs), can be used to infer gene expression changes associated with a given phenotype based on GWAS SNP-effects. This is a powerful approach because association at the vast majority of loci identified through GWAS of complex traits appears to be mediated by altered gene regulation rather than changes in protein coding sequence (6). Several methods exist for inferring associated differential expression from GWAS summary statistics, including Summary-data-based Mendelian randomization (SMR) (7), and transcriptome-wide association study (TWAS, as performed by FUSION (8) and MetaXcan (9)). A key distinction between SMR and TWAS is that TWAS considers the joint effect of multiple SNPs on a gene’s expression and therefore has greater power than SMR when there are multiple eQTLs for a given gene (9,10). In addition to prioritising genes at genome-wide significant loci, TWAS is able to implicate genes in regions containing no genome-wide significant variants. For example, a recent TWAS of schizophrenia identified 157 unique genes as significantly associated, 35 of which were considered as novel as they were >500kb from a genome-wide significant locus (11). Furthermore, by indicating how the regulation of the implicated gene is affected by associated genetic variation, such studies can more accurately inform functional follow-up investigations and, potentially, therapeutic strategies.

In this study, we carry out the first TWAS of ASD in order to identify gene expression changes associated with these disorders. Using the ASD TWAS results, and a novel adaptation of established statistical methods, we also test for enrichment within candidate pathways, specific tissues, and at different stages of brain development. Finally, we estimate the proportion of variance in ASD that is attributable to these TWAS observations.

## Methods and Materials

### Datasets

We performed TWAS using the publically available PGC + iPSYCH ASD GWAS summary statistics (4) (See URLs) and 16 sets of gene expression SNP-weights (Table 1). SNP-weight sets captured gene expression for fetal brain tissue, and brain, blood and adipose tissue in adults. SNP-weights for each gene-tissue pair is referred to as a feature. Fetal brain features were derived using gene expression data collected from brain homogenates from 67 fetuses aged 12-19 weeks post-conception, and genetically-defined to be of European ancestry, collected through the Human Developmental Biology Resource (12). Common mind Consortium (CMC), Netherlands Twin Registry (NTR), Young Finns Study (YFS), Metabolic Syndrome in Men study (METSIM) and Genotype-Tissue Expression project (GTEx) SNP-weights were downloaded directly from the FUSION/TWAS website (see URLs). Information regarding the analysis of genotypes and gene expression from these datasets has been previously described: CMC (11), NTR, YFS, METSIM (8), GTEx (13), fetal brain (12). See also Supplementary Information for further details.

**Table 1:** Descriptive statistics for SNP-weight sets in ASD TWAS.

| Table 1. Descriptive statistics for SNP-weight sets in ASD TWAS. |  |  |  |  |  |
| --- | --- | --- | --- | --- | --- |
| Study | Tissue | Type | N. Individuals | N. Features | ASD TWAS significant |
| O'Brien | Fetal brain | Gene | 67 | 831 | 2 |
| O'Brien | Fetal brain | Transcript | 67 | 2,865 (2,295) | 6 (5) |
| CMC | Dorsolateral prefrontal cortex | Gene | 452 | 5,379 | 1 |
| CMC | Dorsolateral prefrontal cortex | Splicing | 452 | 7,735 (3,297) | 1 |
| NTR | Peripheral blood | Gene | 1247 | 2,437 | 0 |
| YFS | Whole blood | Gene | 1264 | 4,657 | 2 |
| METSIM | Adipose | Gene | 563 | 4,637 | 0 |
| GTEX | Caudate basal ganglia | Gene | 100 | 944 | 0 |
| GTEX | Cerebellar Hemisphere | Gene | 89 | 1,512 | 1 |
| GTEX | Cerebellum | Gene | 103 | 2,001 | 2 |
| GTEX | Cortex | Gene | 96 | 1,047 | 0 |
| GTEX | Frontal Cortex BA9 | Gene | 92 | 928 | 1 |
| GTEX | Hippocampus | Gene | 81 | 539 | 0 |
| GTEX | Hypothalamus | Gene | 81 | 602 | 2 |
| GTEX | Nucleus accumbens basal ganglia | Gene | 93 | 883 | 1 |
| GTEX | Putamen basal ganglia | Gene | 82 | 633 | 0 |
| Total | - | - | - | 37,631 (13,243) | 19 (14) |
| **Note.** *Type*, indicates what the features for each dataset represent i.e. gene-level expression, transcript-level expression, or splicing events; *N. Individuals*, the number of individuals in the reference sample used to derive the feature SNP-weights; *N. Features*, the number of features included in the TWAS for each SNP-weight set. Numbers in parentheses for Fetal brain transcript-level and CMC Dorsolateral prefrontal cortex indicate the number of unique genes. |  |  |  |  |  |

### TWAS

#### Defining transcriptome-wide significance

We estimated transcriptome-wide significance as *p* = 4.25×10^−6^ using a permutation procedure to accurately account for the correlation between features within and across SNP-weight sets (See Supplementary Information).

#### TWAS analysis using FUSION

TWAS analysis was performed using the FUSION software with default settings (see URLs) (8).

Colocalisation was performed to estimate the posterior probability that GWAS and TWAS associations share a causal SNP. Colocalisation was performed using the coloc R package (14), implemented by FUSION. This method uses a Bayesian framework to estimate the posterior probability of five models: Model 0 = No association with either ASD or gene expression, Model 1 = Association with ASD only, Model 2 = Association with gene expression only, Model 3 = Association with ASD and gene expression, but from two independent SNPs, and Model 4 = Association with ASD and gene expression at a common SNP.

In regions containing multiple significant associations, joint analysis was performed to identify conditionally independent associations. This was implemented using FUSION with genes considered in a joint model if the boundaries overlapped ±0.5Mb.

To pool evidence of association for each gene across SNP-weight sets, the multiple degree-of-freedom omnibus test was performed using FUSION.

#### Derivation of non-TWAS informed gene-based statistics

For comparison purposes, MAGMA’s gene-based analyses were performed using the ASD GWAS summary statistics to enable a direct comparison with the TWAS results. This comparison was used to highlight the differences in results between TWAS and a functionally agnostic approach. MAGMA was used to estimate gene-level associations for all unique genes in the TWAS, of which 13,158 contained at least one SNP available in the ASD GWAS and in the 1000 Genomes linkage disequilibrium (LD) reference. The SNP-wise Mean model was used in MAGMA to estimate gene associations, a model also employed by other software (PLINK, VEGAS, SKAT). SNPs were assigned to a gene if they were within 10kb of the gene boundaries.

### TWAS-based enrichment analysis

#### Analytical procedure

TWAS-based enrichment analysis was performed using a novel adaption of a previously established method for GWAS-based enrichment analysis implemented in the software MAGMA (15). In brief, enrichment analysis was performed using linear mixed model regression of TWAS Z-score on gene-set membership, accounting for the correlation between genes due to LD. We analysed TWAS association results from all 16 SNP-weight sets simultaneously to improve genome coverage and reduce the multiple testing burden. The R package lme4qtl was used to fit the linear mixed model (16). The software used for this analysis is publically available (see URLs). See Supplementary Information for further details.

#### Gene-set enrichment analysis

TWAS results were tested for enrichment across 173 candidate gene-sets, including 134 gene-sets relevant to various aspects of nervous system function and development (herein referred to as the central nervous system (CNS) gene-sets), 38 gene-sets that have been previously implicated in ASD specifically (herein referred to as ASD-relevant gene-sets), and a gene-set containing loss of function intolerant genes. The CNS and loss of function gene-sets have been previously described in ref. (17,18), and the ASD-related gene-sets have been previously described in ref. (19). The false-discovery rate (FDR) method was used to correct for multiple testing across all 173 candidate gene sets.

The comparative analysis using non-TWAS informed gene-level associations was also performed using MAGMA.

#### Gene-property association analysis

Gene-property analysis estimates the relationship between TWAS associations and a continuous gene annotation. Using BRAINSPAN data (20), a score indicating preferential expression of each gene at 19 developmental stages has been calculated (11). Using the mixed model approach described above the correlation between preferential expression scores for each developmental period and non-zero association gene Z-scores was then calculated. A significance threshold of *p* < 0.05/19 was used.

For comparison, gene-property analysis was also performed using the non-TWAS informed gene-level associations in MAGMA using default settings.

#### SNP-weight set enrichment analysis

We also tested for an enrichment of association across the SNP-weight sets used in this study to evaluate the importance of each tissue or time point in ASD aetiology. Secondary analysis was also performed using only SNP-weight sets for the basal ganglia, to compare each of the three basal ganglia components to one another.

### Estimating the proportion of heritability mediated by gene expression

The proportions of ASD heritability accounted for by the TWAS results from each SNP-weight set and all SNP-weight sets combined were estimated using stratified-LD score regression (S-LDSC). FUSION was used to calculate LD scores files that were restricted to SNPs within the TWAS SNP-weights and represented the relationship between each SNP and predicted gene expression. The total heritability of ASD was estimated using standard LDSC. The proportion of SNP-based heritability accounted for by TWAS was calculated as the TWAS-based heritability divided by the SNP-based heritability.

## Results

### ASD TWAS

Of the 16 SNP-weight sets, 10 revealed transcriptome-wide significant associations, with the fetal brain transcript-level weights returning the most significant associations (5 unique genes) (Table 1). In total 19 transcriptome-wide significant associations were observed for 14 unique genes (Supplementary Figure 1). Following conditional analysis, 5 independent transcriptome-wide significant associations were observed (Table 2). Many of these associations achieved transcriptome-wide significance across multiple SNP-weight sets (Figure 1). Full TWAS association results are in Supplementary Table 1. Colocalisation posterior probability estimates are available in Supplementary Table 2.

**Table 2.** List of independent transcriptome-wide significant loci.

| Location | MinP<br>(TWAS) | MinP<br>(GWAS) | MinP<br>(MAGMA) | Variance<br>Explained | Jointly<br>significant | Marginally Significant |
| --- | --- | --- | --- | --- | --- | --- |
| chr2:10923518-10952970 | 1.8E-06 | 1.3E-04 | 4.3E-04 | 94.2% | PDIA6 | PDIA6 |
| chr8:8998934-9002945 | 3.3E-07 | 1.8E-06 | 6.6E-07 | 76.5% | RP11-10A14.3 | RP11-10A14.3 |
| chr8:11700033-11725646 | 2.0E-06 | 3.3E-06 | 1.1E-06 | 96.2% | CTSB | CTSB |
| chr17:44344403-44346060 | 5.0E-07 | 4.4E-06 | 1.6E-07 | 99.9% | RP11-259G18.1 | ARHGAP27, CRHR1-IT1,<br>DND1P1, KANSL1,<br>KANSL1-AS1, LRRC37A,<br>LRRC37A4P, MAPT,<br>RN7SL739P, RP11-259G18.1 |
| chr20:21283941-21370463 | 1.8E-08 | 2.0E-09 | 1.9E-09 | 84.5% | XRN2 | XRN2 |

**Note.** Jointly significant, genes that remain significant after accounting for variance explained by all nearby marginally significant genes; Marginally significant, genes that are no longer significant after accounting for variance explained by surrounding jointly significant genes. Associations are considered to be dependent if they are within 1Mb of each other; Location, the chromosome and start/stop coordinates of the jointly significant gene; MinP (TWAS), the minimum p-value across SNP-weight sets for the jointly significant gene; MinP (GWAS), indicates the p-value for top SNP-association +/-500kb of the jointly significant gene; MinP (MAGMA), p-value of most significant gene in MAGMA analysis +/-500kb of the jointly significant gene; Variance explained, the proportion of the MinP (GWAS) association explained by the most significant TWAS feature in the region, calculated as $1 - (\chi^2_{\text{conditioned GWAS association}} / \chi^2_{\text{unconditioned GWAS association}})$ .

**Figure 1.**
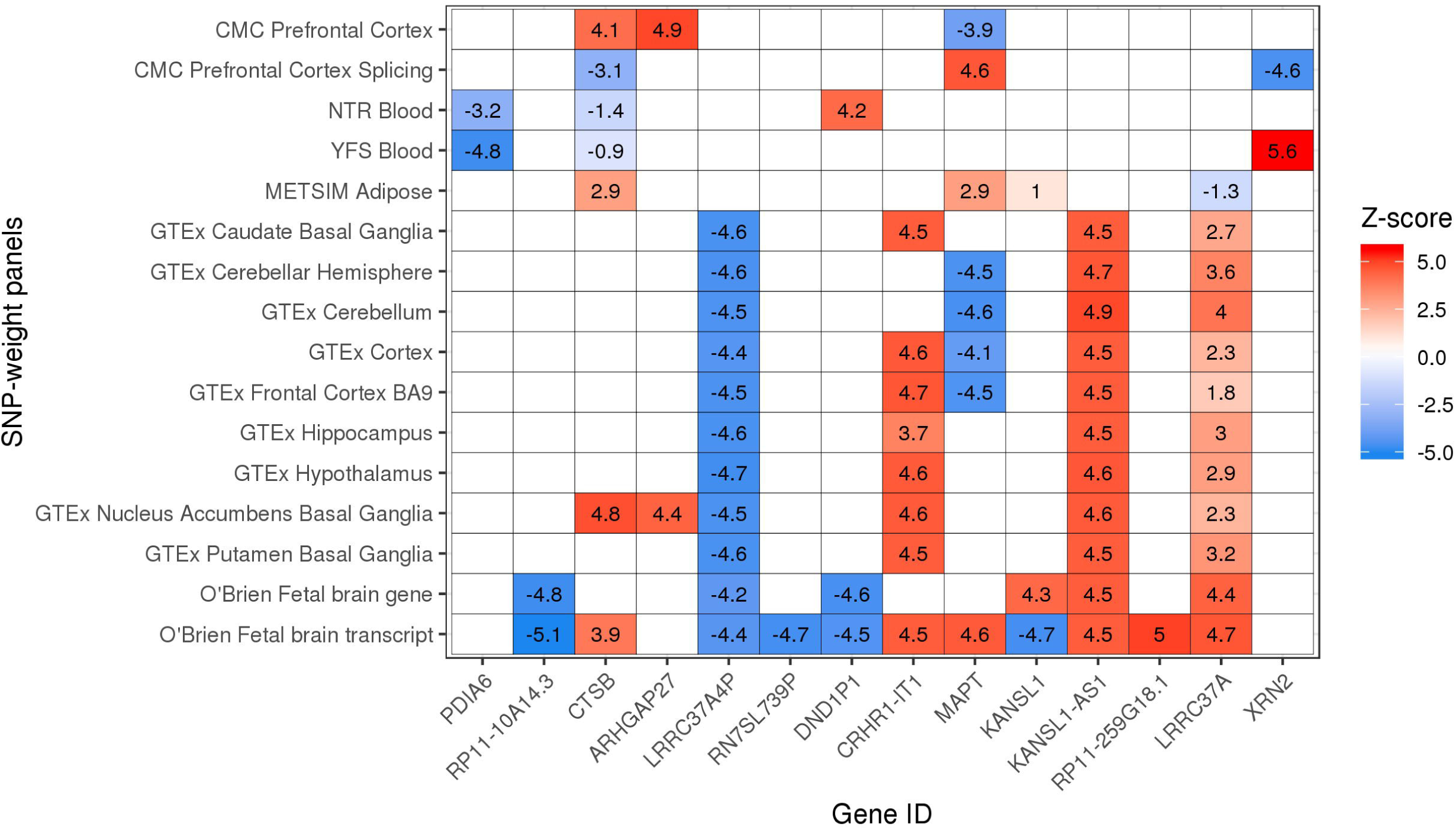
Transcriptome-wide significant genes across SNP-weight sets. Transcriptome-wide significance as a Z score is ∼4.6. Note. The direction of effect for splicing and transcript SNP-weights should be interpreted with caution. Blank squares indicate the gene weights were not available in the target tissue.

#### chr20 p11.22

The strongest ASD TWAS association was the upregulation of *XRN2* based on YFS blood SNP-weights (*p* = 1.80×10^−8^). Differential splicing of *XRN2* also showed suggestive significance based on the CMC prefrontal cortex SNP-weights (*p* = 4.86×10^−6^), and as a result the omnibus test *p*-value for *XRN2* was 1.50×10^−8^. Colocalisation analysis supports Model 4 with a posterior probability of 0.966, providing evidence that ASD liability and *XRN2* expression are causally associated with the same variant. *XRN2* is within a locus previously associated with ASD at genome-wide significance in the ASD GWAS, with predicted expression of *XRN2* explaining 84.5% of the top SNP association (Table 2, Figure 2).

**Figure 2.**
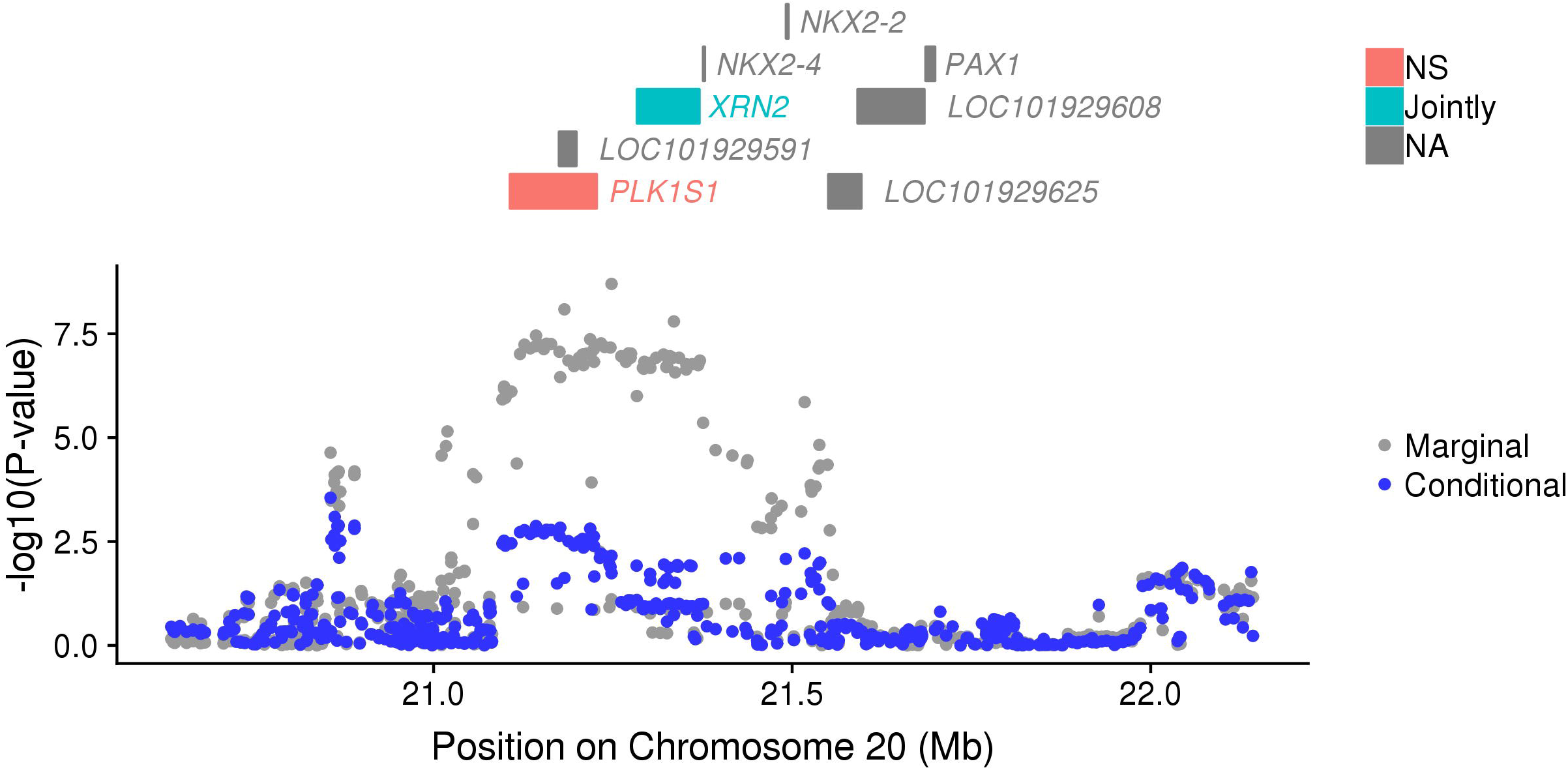
Regional association plot. The top panel shows all of the protein-coding genes or genes in the TWAS. Jointly significant genes are highlighted in blue, non-significant genes are highlighted in red, and genes that were not in the TWAS are in grey. The bottom panel shows a Manhattan plot of the GWAS data before (gray) and after (blue) conditioning on the jointly significant genes.

#### chr17 q21.31

A cluster of 14 transcriptome-wide significant associations (10 unique genes) were observed within a 1Mb region on chromosome 17 corresponding to an inversion polymorphism that is common in European populations (21). No single SNP within this region achieved genome-wide significance in the ASD GWAS. The most significant TWAS association in this region was the upregulation of an *RP11-259G18.1* transcript in the fetal brain, explaining 99.9% of the ASD SNP association in this region (Supplementary Figure 2). Features in this region were highly correlated (Supplementary Figure 3) and therefore represent a single association. Although *RP11-259G18.1* showed the strongest TWAS association, colocalisation analysis supported Model 4 (same causal variant as ASD) for all transcriptome-wide significant associations in this region.

**Figure 3.**
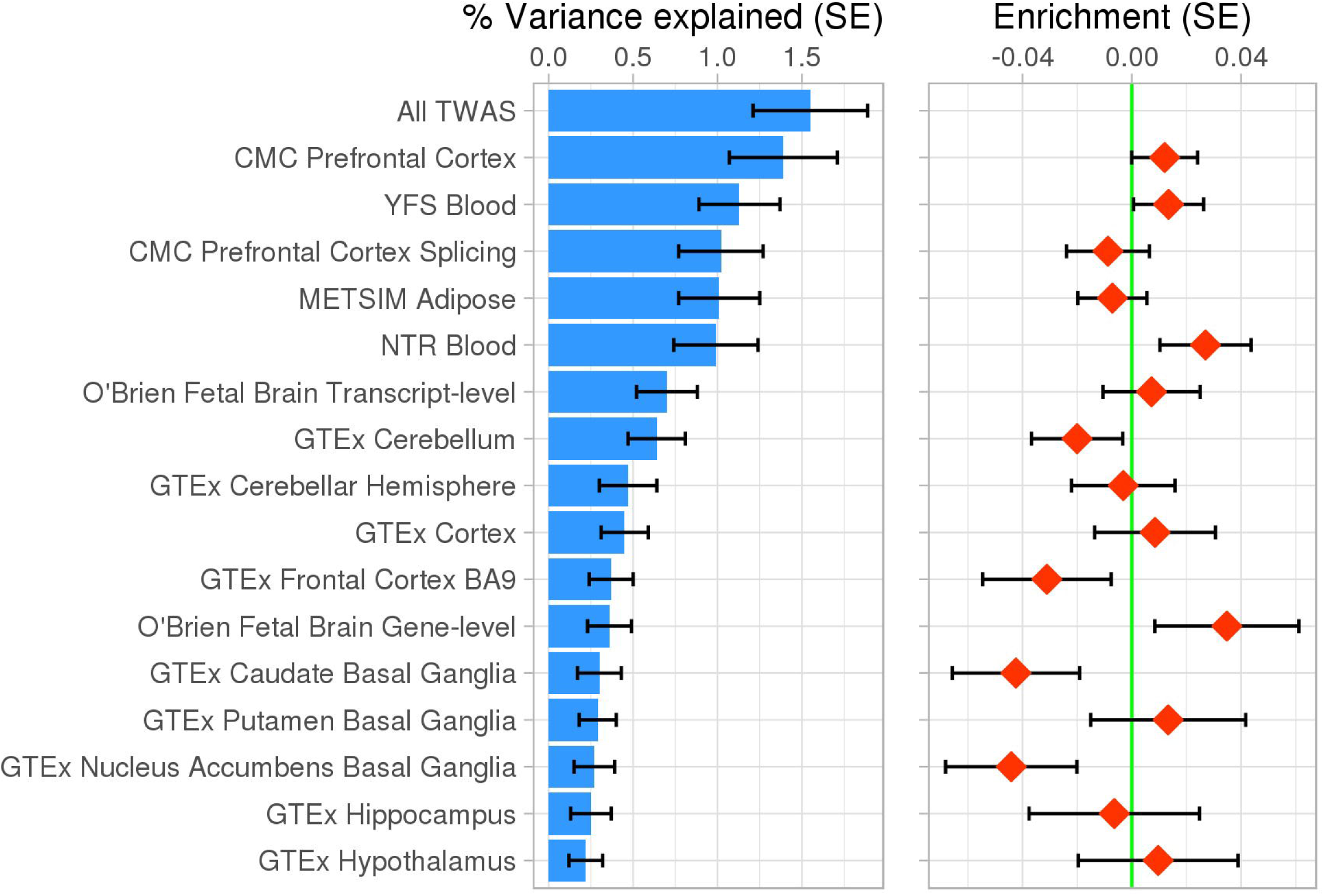
The left panel shows the ASD SNP-heritability explained by predicted gene expression on a liability scale. The right panel shows results of competitive gene set enrichment analysis for SNP-weight sets (i.e. whether features within each SNP-weight set are on average more associated with ASD than compared to features in all other SNP-weight sets.

#### chr8 p23.1

Two genes, *CTSB (*nucleus accumbens basal ganglia) and *RP11-10A14.3* (fetal brain gene- and transcript-level), located 2.7Mbs apart on chromosome 8 achieved transcriptome-wide significance. When considered in a joint model, both the transcript-level feature for *RP11-10A14.3* and *CTSB* remained nominally significant, indicating that the signal driving the ASD associations with *RP11-10A14.3* and *CTSB* are broadly independent. Although there are no genome-wide significant SNPs within 500kb of these genes (the default criteria for being ‘novel’ (11)), they are either side of a genome-wide significant locus (rs4841432, chr8:10583506, *p*=4.4×10^−8^) (Supplementary Figure 4). In the joint model, the expression of *CTSB* and *RP11-10A14.3* together explain 60% of the association for this genome-wide significant SNP, demonstrating that these TWAS associations are correlated with the previously identified genome-wide significant association, and are therefore not entirely novel. Colocalisation provides weak evidence that these association a driven by the same causal variant as ASD as the posterior probability is greater for Model 4 (same causal variant) than Model 3 (different causal variant), but individually weak SNP effects result in other models being the preferred model (Supplementary Table 2).

#### chr2 p25.1

The transcriptome-wide significant association between *PDIA6* and ASD on chromosome 2 is in a genomic region showing minimal evidence of association at the SNP-level, with the minimum *p*-value being 1.3×10^−4^ (+/-2Mbs of *PDIA6*) (Supplementary Figure 5). The data best supports Model 4 in which ASD and PDIA6 share a single causal association.

### Gene set and property analysis

Full competitive gene-set enrichment results are available in Supplementary Table 3. Of the 135 candidate gene sets, 15 achieved nominal significance, with the most significant being ‘Synaptic vesicle’, ‘Presynapse’ and ‘Abnormal axon guidance’ (*p* < 0.015). No gene sets were significant after FDR correction. Of the 38 ASD candidate sets, one achieved nominal significance (called ‘Parikshak2013_M16’) with none achieving significance after FDR correction. The M16 coexpression module represents early cortical development, with upregulation of this module starting at 10 weeks post conception (22). The loss-of-function intolerant gene-set returned an enrichment *p*-value of 0.194, which supports the notion that mutation-intolerance metrics do not characterise ASD GWAS loci despite their association with ASD risk genes identified through *de novo* variant studies (23).

Enrichment results of genes preferentially expressed during 1 of 19 developmental periods in brain, returned no associations achieving nominal significance (Supplementary Figure 6). However, preferential expression during 7 of the first 8 developmental stages (12PCW – 4 months) were positively correlated with ASD TWAS association, and 7 out of 11 later stages (10 months – 40 years) were negatively correlated with ASD TWAS association. This trend suggests that ASD TWAS associations are relatively enriched among genes showing high expression during fetal development.

Enrichment analysis comparing the mean association of features within each SNP-weight set showed no significant enrichment (Figure 3), with fetal gene expression showing the highest level of enrichment (*p* = 0.09). Secondary competitive analysis of the three basal ganglia regions alone showed that gene expression in the putamen region was enriched at nominal significance in comparison to gene expression in the caudate and nucleus accumbens (*p* = 0.03).

### Comparison of TWAS results to MAGMA

MAGMA gene association analysis returned similar results to those reported previously by Grove and colleagues (4). Regions containing transcriptome-wide significant associations on chromosome 8, 17 and 20 also contained significant MAGMA-based associations (±500kb of significant TWAS feature), although the different methods often implicated different genes within the same locus (Supplementary Tables 4 and 5, Supplementary Figures 7-12). The only transcriptome-wide significant locus in which MAGMA identified no significant associations was that surrounding *PDIA6* on chromosome 2. MAGMA identified three loci containing significant genes that contained no TWAS significant genes.

Similar to TWAS-based gene set enrichment analysis, MAGMA-based gene set analysis using ASD GWAS summary statistics of candidate gene sets returned no significant associations after FDR correction (Supplementary Tables 3). The rank-based correlation between MAGMA- and TWAS-based gene set association test statistics was 0.23.

Gene property analysis for enrichment of genes preferentially expressed during a given period of brain development, showed a similar pattern of results as the TWAS-based gene property analysis, with a rank-based correlation of 0.39. Although, no developmental stage achieved significance in the MAGMA analysis after Bonferroni correction, preferential expression in 3 fetal stages of brain development were positively associated at nominal significance, and preferential expression in 1 adult brain stage (>19 years) was negatively associated at nominal significance (Supplementary Figure 6).

### Proportion of heritability

LDSC estimated the total ASD SNP-heritability at .120 (SE = .010, *p* = 4.72×10^−32^) on a liability scale assuming a population prevalence of 1.2%. When considering the TWAS results from all SNP-weight sets together, the heritability was .0155 on a liability scale, and the proportion of ASD SNP-heritability explained was 13%. The TWAS-based heritability estimates were all significantly non-zero, with the proportion of heritability explained by each TWAS showing a positive correlation with both the number of features available and the number of individuals used to derive the SNP-weights (Figure 3, Supplementary Table 6).

## Discussion

This is the first study to infer differential gene expression/splicing associated with ASD using the TWAS method and has provided several novel insights into the aetiology of ASD.

This study has demonstrated that the previously reported genome-wide significant locus spanning multiple genes within the locus at 20-21 Mb of chromosome 20 is linked to significant differential expression and splicing of the gene *XRN2*. Functionally agnostic gene-based analysis in MAGMA also identifies *XRN2* as significant, as reported in this study and previously by Grove and colleagues (4). Moreover, a recent study reported evidence that ASD associated SNPs in this region colocalise with several DNA methylation sites (24), although the consequence of this methylation on surrounding gene expression is unknown. Our data point to differential expression of *XRN2* in the blood and differential splicing in the prefrontal cortex. *XRN2* is an essential nuclear 5’→3’ exoRNase with a multitude of functions in the processing and regulation of RNA molecules. *XRN2* has been identified as an essential gene for the survival of multiple human cell lines (25–27), and individuals diagnosed with ASD have been shown to have an increased number of deleterious mutations among essential genes (28), again supporting a role for *XRN2* in ASD aetiology.

This study also highlighted 13 transcriptome-wide significant genes outside loci achieving genome-wide significance in the corresponding ASD GWAS (±500kb), 10 of which surround the 17q21 inversion. Five of the 10 significant genes surrounding the 17q21 inversion are also identified as significantly associated using the functionally agnostic region-based approach employed by MAGMA, of which several were previously reported by Grove and colleagues (4). The inversion at 17q21, which has a population frequency of around 20% in Europeans, has been previously highlighted as an ASD susceptibility locus through linkage analysis (29) and family-based GWAS (30). The 900kb inverted region, which contains many known genes, is marked by extensive LD, complicating identification of the causal susceptibility genes. One study used a fine mapping approach and implicated *CACNA1G* as a ASD susceptibility gene in the region (31). Results from our TWAS show no evidence of association between ASD and *CACNA1G* expression (*p* = 0.33 based on CMC Prefontal Cortex). Expression of several other genes mapping to this region have recently been implicated in the personality trait of neuroticism (12).

The only locus containing a transcriptome-wide significant gene that is not significantly implicated by either GWAS or MAGMA was *PDIA6* on chromosome 2. This discovery highlights the advantage of TWAS which incorporates additional functional information of genetic variants as opposed to relying purely on the proximity of SNPs to a gene.

Previous studies using the TWAS approach commonly report associations as novel if the associated feature is outside of genome-wide significant loci in the corresponding GWAS (11,32). However, given that TWAS is a gene-based approach, it gains power both from incorporating functional annotations but also by pooling evidence across multiple genetic variants. Therefore, we have compared the TWAS results to those from the functionally agnostic gene-based approach employed in MAGMA (and other software) to more clearly distinguish the novel insights that TWAS can provide. This comparison demonstrated that four of the five regions containing independent ASD TWAS associations also contained significant associations identified by the functionally agnostic approach, suggesting that it is often pooling information across genetic variants that highlights regions of novel association. However, the genes that were implicated within these regions often differed between the two gene-based approaches. A key advantage of TWAS is that it considers the functional annotations of associated genetic variants and can therefore provide mechanistic insight into how a regional association is mediated. This is valuable information for subsequent experimental studies that aim to understand the mechanism underlying the genetic association, and could also be used to improve subsequent gene-level statistical analyses. However, TWAS only assesses genes that show statistically significant cis-heritable expression, and is therefore dependent on the sample size of the gene expression reference. As functionally agnostic region-based approaches do not suffer from this limitation, we consider the two approaches complimentary, with TWAS as a useful downstream approach for refining and assigning directionality to gene associations.

The two transcriptome-wide significant associations on chromosome 8, *CTSB* and *RP11-10A14.3,* were proximal to a genome-wide significant locus in the corresponding GWAS. MAGMA analysis identified six genes within this region achieving significance (Supplementary Tables 4 and 5), however no gene was identified as significant by both TWAS and MAGMA. Additional support for this locus comes from repeated studies showing duplications in this region (8p23.1-3) in individuals with ASD (33–35). However, the gene/genes driving this association have not been identified. This TWAS found evidence that *CTSB* is upregulated in ASD across multiple brain tissues. *CTSB* encodes Cathepsin B, a cysteine protease that has been reported as a mediator of exercise enhanced hippocampal neurogenesis and spatial memory (36), and inhibitors of Cathepsin B have therapeutic potential for traumatic brain injury (37). Furthermore, treating rodent neuroprogenitor cells with exogenous Cathepsin B is associated with differential expression of multiple neurogenesis-related genes (36). These previous findings suggest that differential expression of *CTSB* leads to several differences in neurogenesis and neuronal cell death, and therefore is a plausible candidate for ASD. Cathepsin B is also an amyloid precursor protein secretase, and inhibition of it has been reported as a potential therapeutic for Alzheimer’s disease (38). This is interesting given prior evidence of shared aetiology between ASD and Alzheimer’s disease (39). *RP11-10A14.3* is an antisense RNA with an unknown function. Several other genes in this region show suggestive evidence of differential expression/splicing in ASD (Supplementary Figure 8), including *MSRA*, which has been previously associated with schizophrenia (40), *MFHAS1*, and *PINX1*.

Finally, of the TWAS implicated genes, downregulation of *PDIA6* in the blood was significantly associated with ASD. *PDIA6* encodes a member of the protein disulphide isomerase (PDI) family, which play an important role in protein folding. PDIs are important for forming, breaking and rearranging disulphide bonds, and as general chaperones. As a results of their role in protein folding, they have been implicated in a number of neurodegenerative diseases (41), however there is little evidence of a connection between PDIs and neurodevelopmental phenotypes. Further research into the potential role of PDI proteins in ASD is needed.

Gene set and property enrichment based on TWAS associations showed limited success, with several interesting observations, but none surviving multiple testing correction. Analogous gene-level analyses using MAGMA also failed to identify significant associations, indicating that the null findings could be a consequence of the as yet low power of the ASD GWAS, rather than limitations of the TWAS-based enrichment analysis approach. Further work exploring the utility of TWAS associations in enrichment analyses is warranted.

Predicted gene expression based on all SNP-weights sets separately and together explained a significant amount of variance in ASD liability, collectively accounting for 13% of the ASD SNP-heritability. This supports the notion that TWAS is a useful approach for understanding the aetiology of ASD, although this estimate may be upward biased as it captures heritability explained by predicted expression and all heritable variation that is correlated with the predicted expression.

There are two key limitations to this study. Firstly, TWAS identifies genetic variation which is associated with two outcomes (in this case ASD and gene expression/splicing), with subsequent colocalisation analysis to determine whether the association is driven by linkage (two causal SNPs in LD with each other) or pleiotropy (same causal SNP). However, neither TWAS nor colocalisation can determine whether the association is causal (the expression mediates the association between SNP and phenotype). Additional studies are required to validate the causal relationship between gene expression changes and ASD. Secondly, the SNP-weights used for predicting differential expression/splicing are based on relatively small sample sizes and therefore cannot infer all features that are cis-regulated across all tissues. As a consequence, there may be features that are important for the aetiology of ASD that we are unable to capture currently using TWAS. Looking forward, more accurate predictions of gene expression afforded by larger expression QTL studies will improve the power of TWAS for identifying those gene expression changes relevant to complex traits and disorders.

This study has provided several insights into the genetic basis of ASD through inference of differential gene expression associated with ASD based on the latest ASD GWAS summary statistics and summary-level gene expression data from multiple tissues, including fetal and adult brain. This study has highlighted differences between TWAS and a common functionally agnostic gene-based approach, and developed a novel procedure for TWAS-based enrichment analysis.

## Supporting information

## Acknowledgements

We would like to thank Alexander Gusev for advice on the use of FUSION, and for sharing the preferential expression score based on BRAINSPAN data. We would also like to thank the PGC ASD Working Group for making the ASD GWAS results publically available. This work was supported by an MRC Centre grant (MR/L010305/1), and Medical Research Council (UK) project grant to NJB (MR/L010674/2).

## URLs

ASD GWAS Summary statistics - https://www.med.unc.edu/pgc/results-and-downloads

FUSION software and SNP-weights based on CMC, NTR, YFS, METSIM, and GTEx datasets - http://gusevlab.org/projects/fusion/

TWAS-based enrichment software - https://github.com/opain/TWAS-GSEA

PLINK - https://www.cog-genomics.org/plink/1.9/

