## Supplementary material for "Novel Insight into the Aetiology of Autism Spectrum Disorder Gained by Integrating Expression Data with Genome-wide Association Statistics"

Pain et al.

### Supplementary Text

#### Datasets

##### PGC + iPSYCH ASD GWAS summary statistics

The PGC + iPSYCH ASD GWAS summary statistics are publicly available and were downloaded from: <https://www.med.unc.edu/pgc/results-and-downloads>. The downloaded ASD GWAS summary statistics contained 7,583,402 variants. Additional quality control and processing was performed, including nomenclature and genome build synchronised to the 1000 genomes reference dataset, exclusion of insertion/deletions, and exclusion of strand ambiguous and variants with an INFO score of less than 0.8. A final dataset of 6,038,890 single nucleotide polymorphisms (SNPs) was carried forward for analysis.

*Gene Expression Datasets*

The 16 expression SNP-weight sets are listed in Table 1, including gene expression SNP-weights for fetal brain tissue, and brain, blood and adipose tissue in adults. Features were derived using data collected from the dorsolateral prefrontal cortex of 621 individuals collected by the CommonMind Consortium (CMC) (1), peripheral blood from 1,245 individuals from the Netherlands Twin Registry (NTR) (2), blood from 1,264 individuals within the Young Finns Study (YFS) (3, 4), adipose tissue from 563 individuals from the Metabolic Syndrome in Men study (METSIM) (5), 9 specific brain regions in 81-103 individuals in the Genotype-Tissue Expression project (GTEx) (6), and brain homogenates from 67 genetically-determined European fetuses aged 12-19 weeks post-conception which were collected through the Human Developmental Biology Resource (HDBR) (7). Features used in this TWAS primarily represent gene expression, but additional transcript specific expression features are available in the O’Brien fetal brain dataset, and features representing splice events are available in the CMC dorsolateral prefrontal cortex dataset.

CMC, NTR, YFS, METSIM and GTEx SNP-weights were downloaded directly from the FUSION/TWAS website (see URLs). Information regarding the analysis of genotypes and gene expression from these datasets has been previously described previously: CMC (8), NTR, YFS, METSIM (9), GTEx (6), O’Brien fetal brain (7).

From the processed genotype and gene expression data, TWAS predictors were computed using FUSION software (see URLs).

#### TWAS

##### Defining transcriptome-wide significance

Many genes are available in multiple SNP-weight sets and have highly correlated predicted expression. Furthermore, genes near one another often have correlated expression and are therefore not independent. Therefore, a Bonferroni significance threshold for all features (gene/tissue combinations) would be highly conservative. We estimated transcriptome-wide significance using a permutation procedure, which accounts for the correlation between features, within and across SNP-weight sets. Initially, expression levels for all features from all SNP-weight sets (N features = 38,157) were imputed into the 1000 genomes reference dataset used by FUSION (N individuals = 489) using the FUSION protocol which involves PLINK (10) (see URLs). For each permutation, a random normally distributed phenotype was generated, linear regression was performed to derive a *p*-value of association for each feature, and the minimum *p*-value was stored. This procedure was repeated 1,000 times. The 5% quantile of the minimum p–values is the transcriptome-wide significance threshold with the features used in this study. Based on these permutation the transcriptome-wide significance threshold was estimated at *p* = 4.25×10^-6^ (95% CI = 2.86×10^-6^ – 5.53×10^-6^).

##### Calculating proportion of SNP association explained by predicted expression

The proportion of a SNP-level association accounted for by predicted expression in the TWAS was calculated as 1-(χ2 of conditioned GWAS association) / χ2 of unconditioned GWAS association). This is the same method used by TWAS-hub to calculate ‘% variance explained’ (http://twas-hub.org).

#### TWAS-based enrichment analysis

##### Analytical procedure

Competitive enrichment was tested for by performing a linear regression for each gene-set, whereby gene Z-scores were predicted by membership of each gene-set, including covariates for gene length and the number SNPs within the gene region. Given the functional consequence of each genes up- or down-regulation is unknown, Z-scores were calculated as probit transformed (1-*p*), resulting in an approximately normally distributed Z-score of non-zero association. To avoid potential bias due to outliers, Z-scores were truncated to be between -3 and 6. This regression approach for enrichment analysis can also be used to test for a correlation between TWAS associations and continuous gene annotations, termed gene property analysis, as is also implemented in MAGMA.

To avoid bias due to the correlation between genes we use lme4qtl (11) to fit a mixed model regression of TWAS Z-score on gene-set membership, accounting for the correlation in Z-scores between genes due to LD. The correlation matrix used was computed based on the same predicted gene expression values used when estimating the transcriptome wide-significance threshold. The correlation between genes that were more than 5Mb apart or on separate chromosomes were set to zero. Any gene-gene correlations with an R-squared less than 0.0001 were set to zero. The matrix was stored as a sparse matrix substantially reducing the memory requirements and duration of the analysis. The linear mixed model with the sparse matrix was performed using the lme4qtl package in R. The software used for this analysis (TWAS-GSEA) is publically available (see URLs).

We analysed TWAS association results from all 16 SNP-weight sets simultaneously to improve genome coverage and reduce the multiple testing burden. If multiple features represent the same gene, such as when a gene is captured in multiple SNP-weight sets, only the feature that gave the best prediction of expression (as measured by cross-validated R2) was retained.

For gene set analysis, the feature IDs were converted to entrez IDs using the biomaRt package in R, matching based on the ‘external_gene_name’ variable. Of the unique TWAS features, 11,470 had entrez IDs and could be included in the analysis. The MAGMA gene-set analysis was performed using the MAGMA derived gene-level associations and default settings. 11,424 genes in the MAGMA analysis had entrez IDs available and could be included in the gene-set analyses.

For gene property analysis, of the unique TWAS features, 8,699 were present in the BRAINSPAN preferential expression dataset. For comparative gene property analysis in MAGMA, 8,685 genes in the MAGMA gene analysis were present in the BRAINSPAN preferential expression dataset and were included in the analysis.

### Supplementary Figures

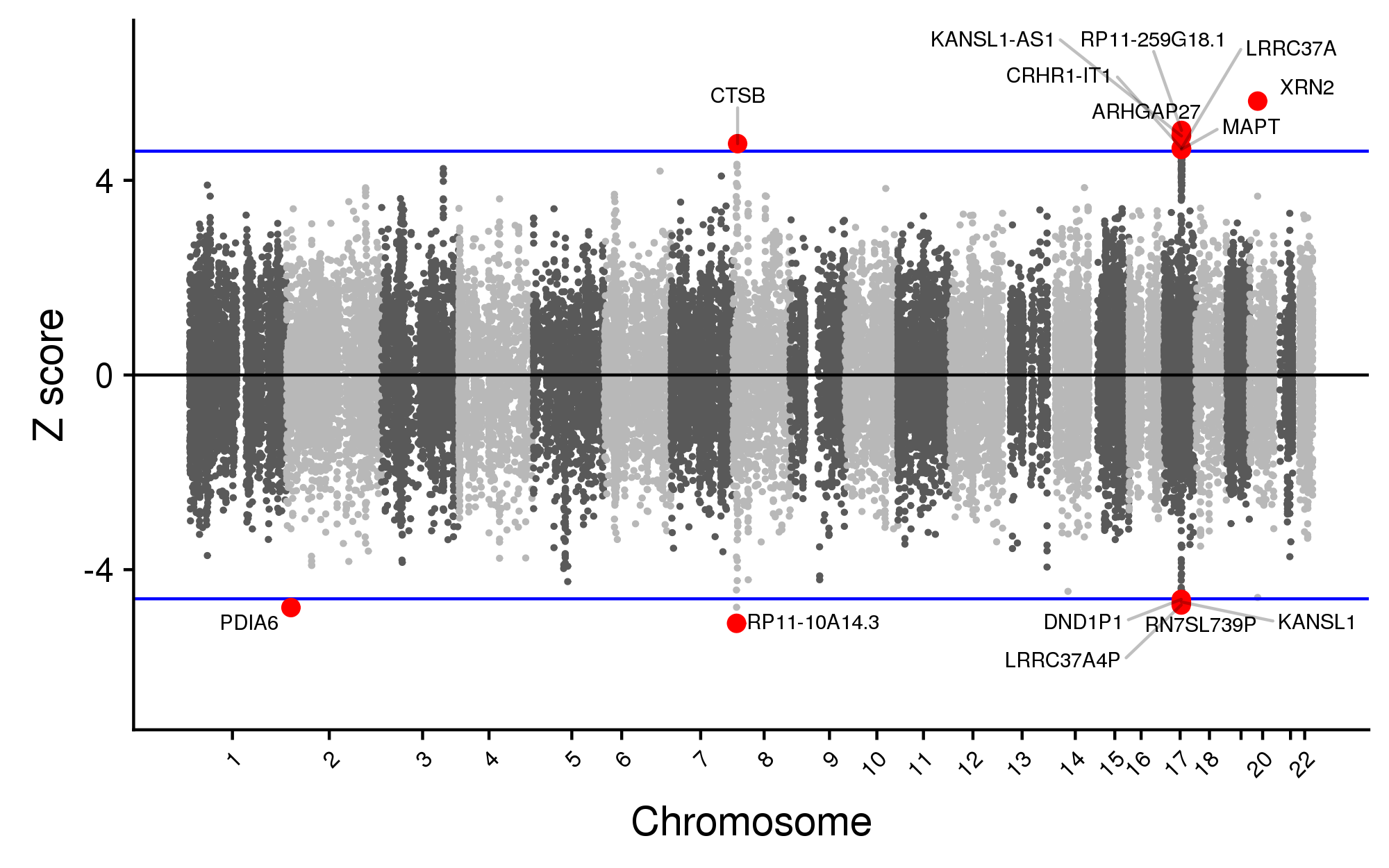

Supplementary Figure 1. Manhattan plot of all ASD TWAS associations. Each point represents a single gene tested, with physical position plotted on the x-axis and Z score of association between the gene and ASD plotted on the y-axis. Transcriptome-wide significant associations are highlighted as red points and are labelled with their ID. If more than one transcriptome-wide significant feature represents the same gene, only the most significant feature is highlighted in red and labelled. The blue horizontal line indicates transcriptome-wide significance (p < 4.25×10^-6^).

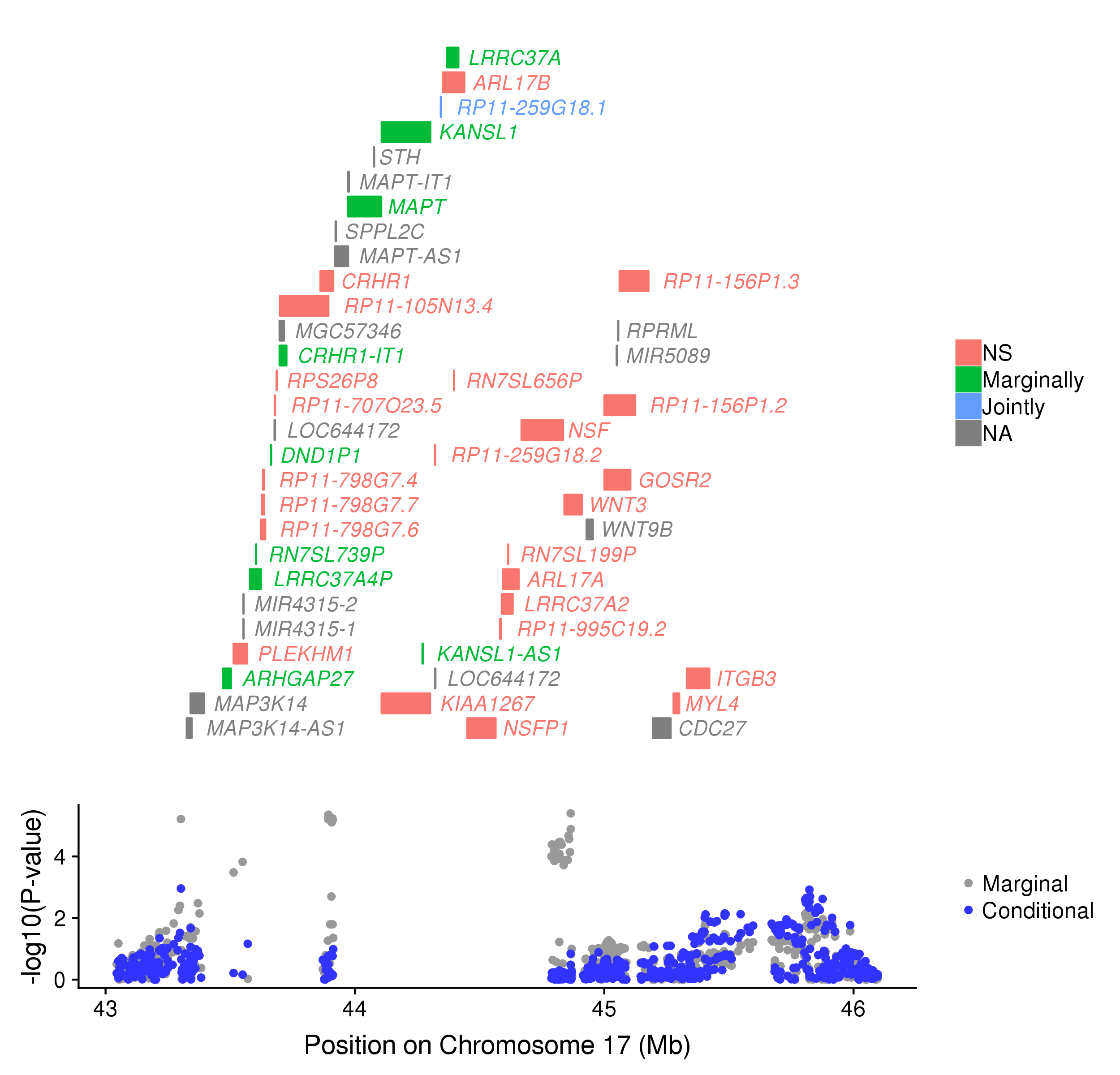
Supplementary Figure 2. Regional association plot. The top panel shows all of the genes in the locus. Marginally TWAS associated genes are highlighted in green, jointly significant genes are highlighted in blue, non-significant genes are in red, and genes that were not assessed in the TWAS are in grey. The bottom panel shows a Manhattan plot of the GWAS data before (gray) and after (blue) conditioning on the green genes.

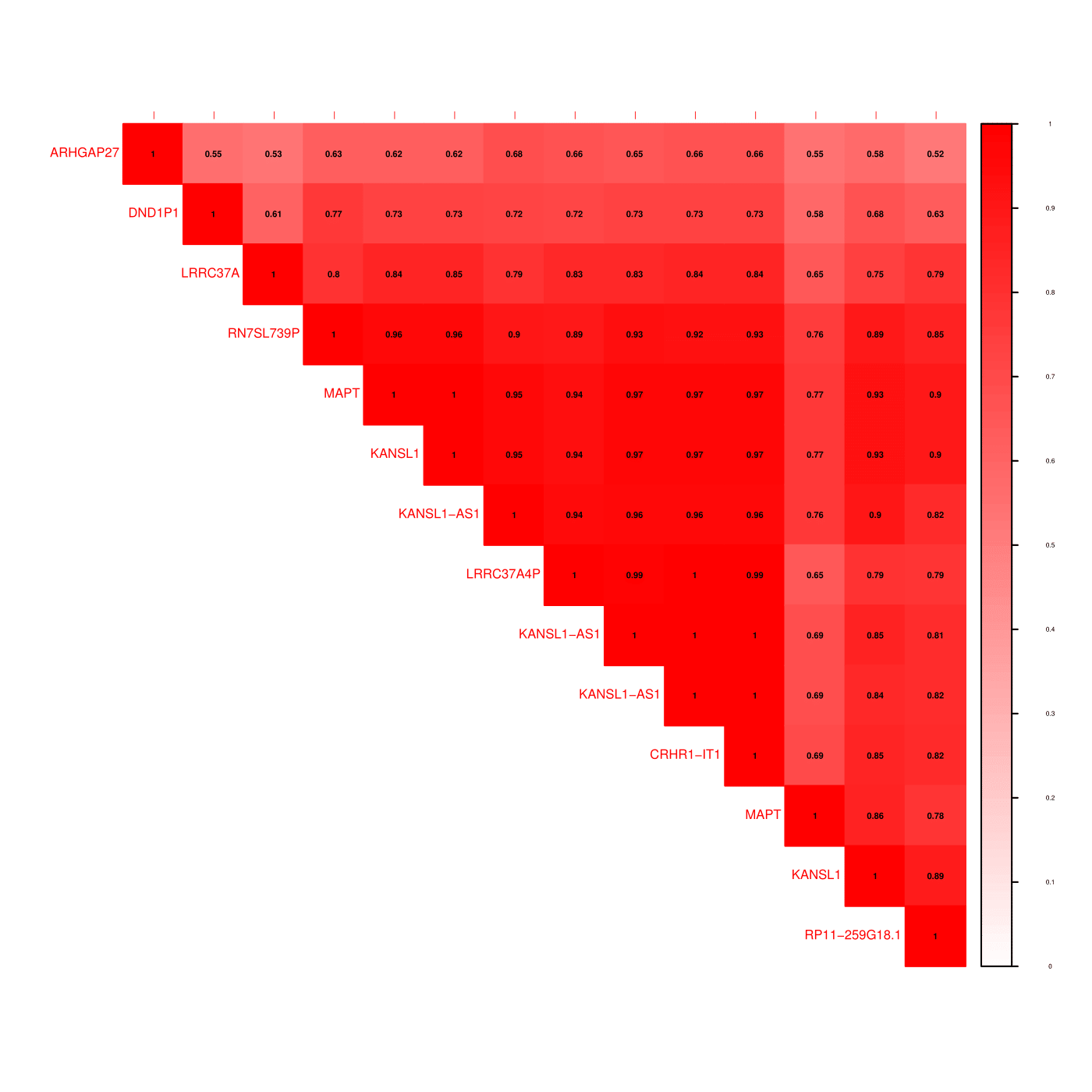

Supplementary Figure 3. Correlations between transcriptome-wide significant genes on chromosome 17.

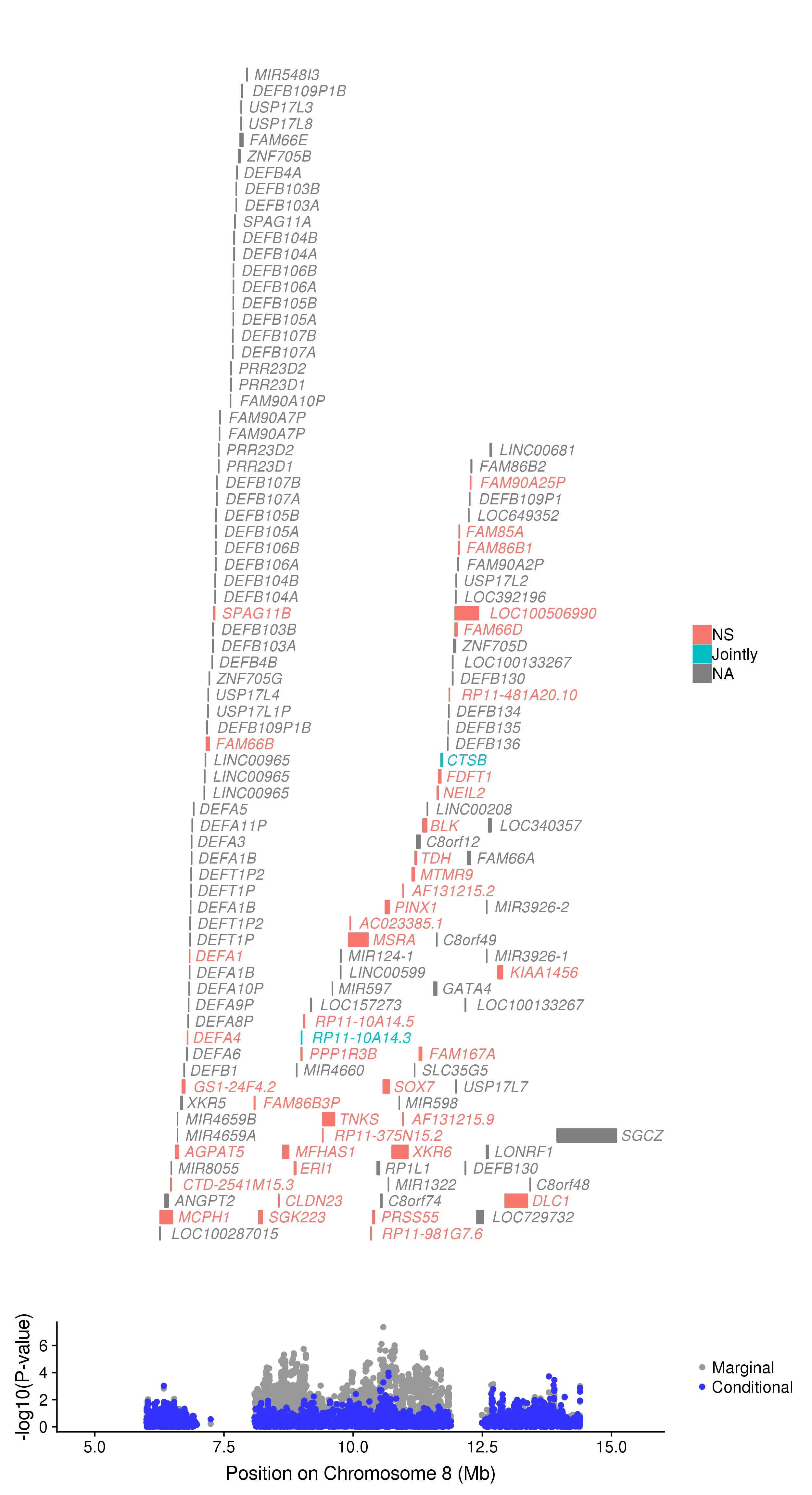

Supplementary Figure 4. Regional association plot. The top panel shows all of the genes in the locus. Marginally TWAS associated genes are highlighted in green, jointly significant genes are highlighted in blue, non-significant genes are in red, and genes that were not assessed in the TWAS are in grey. The bottom panel shows a Manhattan plot of the GWAS data before (gray) and after (blue) conditioning on the green genes.

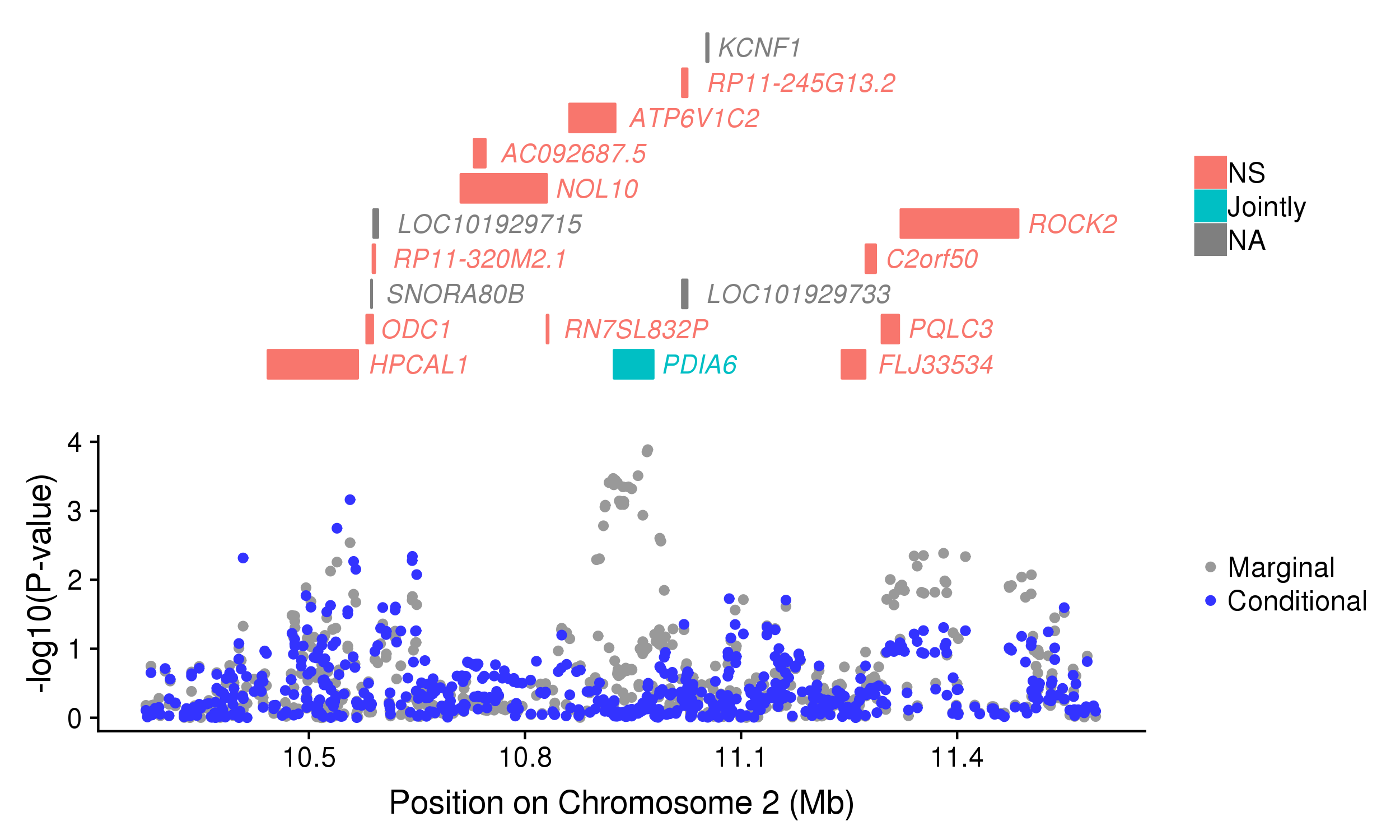
Supplementary Figure 5. Regional association plot. The top panel shows all of the genes in the locus. Marginally TWAS associated genes are highlighted in green, jointly significant genes are highlighted in blue, non-significant genes are in red, and genes that were not assessed in the TWAS are in grey. The bottom panel shows a Manhattan plot of the GWAS data before (gray) and after (blue) conditioning on the green genes.

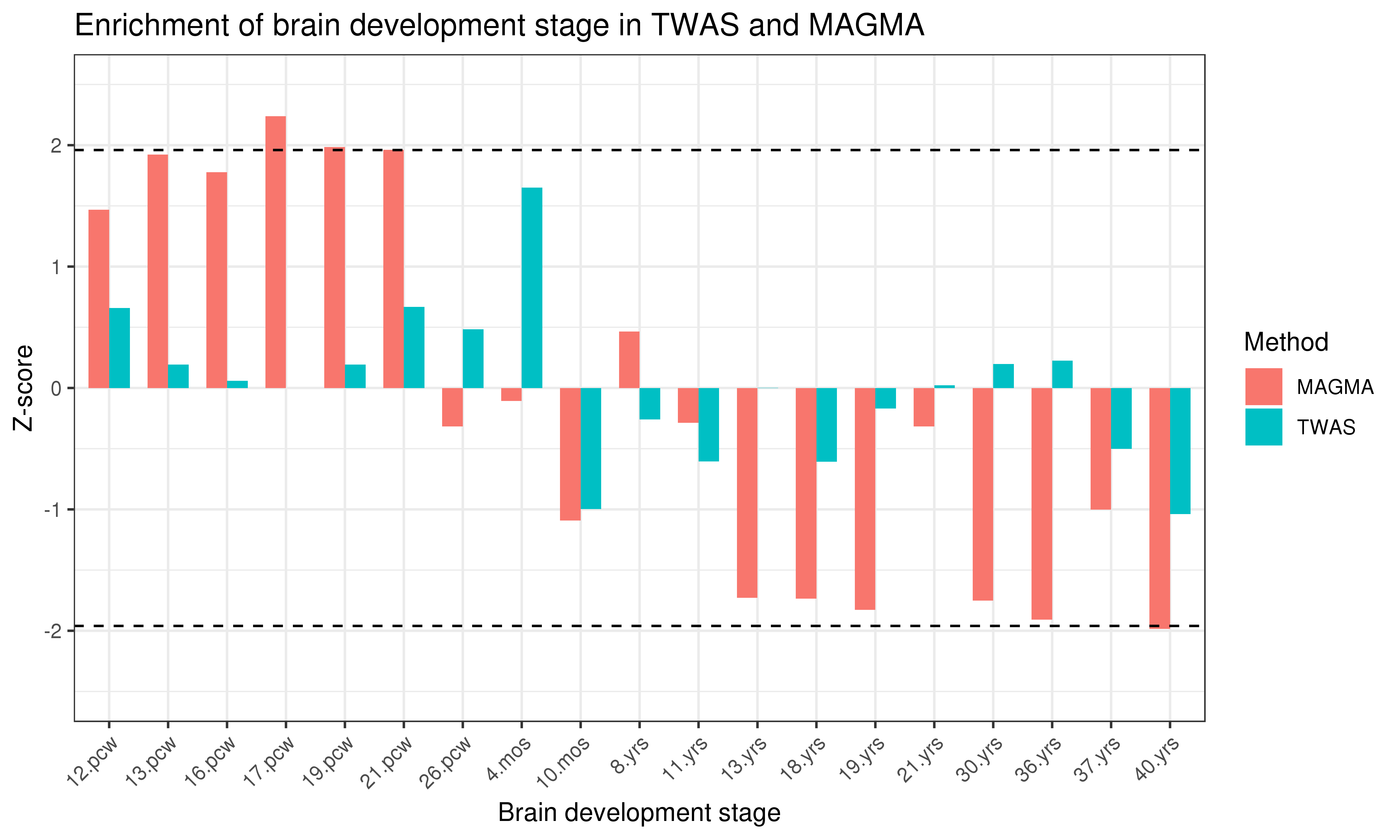

Supplementary Figure 6. Gene property analysis of preferential gene expression across brain development. Results based on TWAS derived Z scores and MAGMA derived Z-scores are shown. The dashed line indicate nominal significance (p=0.05).

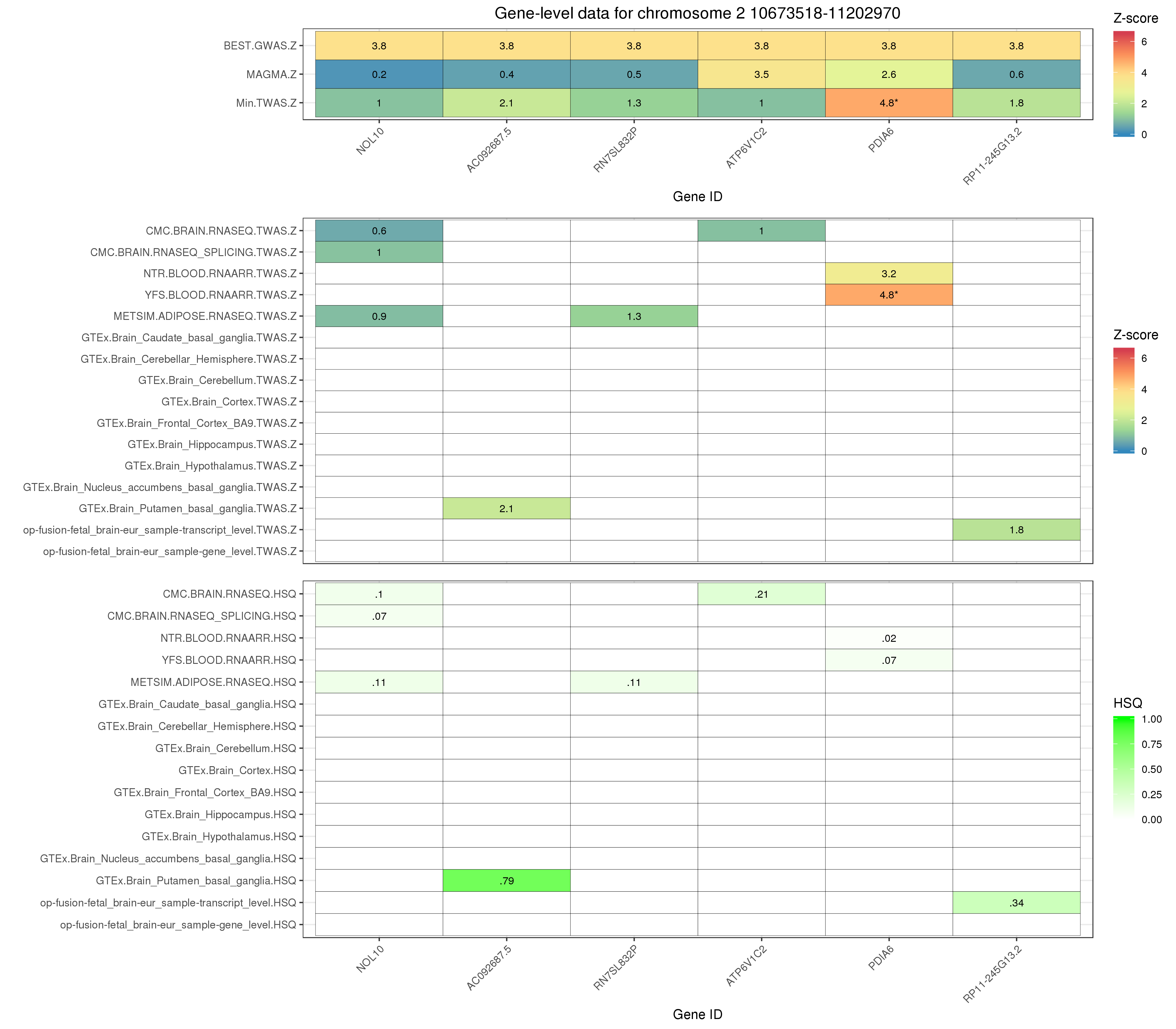

Supplementary Figure 7. Comparison of gene-level Z scores derived using MAGMA and TWAS, and SNP-level Z score in the corresponding GWAS, containing either a significant TWAS or MAGMA association. The absolute TWAS Z score is used here. Genes within 250kb of genes significant in the either the TWAS of MAGMA analysis are included. Empty cells indicate the gene was not available in the TWAS or MAGMA analysis. Asterisks indicate the Z score surpassed the corresponding significance threshold (TWAS = p < 4.25x10^-6^; MAGMA = Bonferroni p-value </= 0.05, GWAS = p <=5x10^-8^).

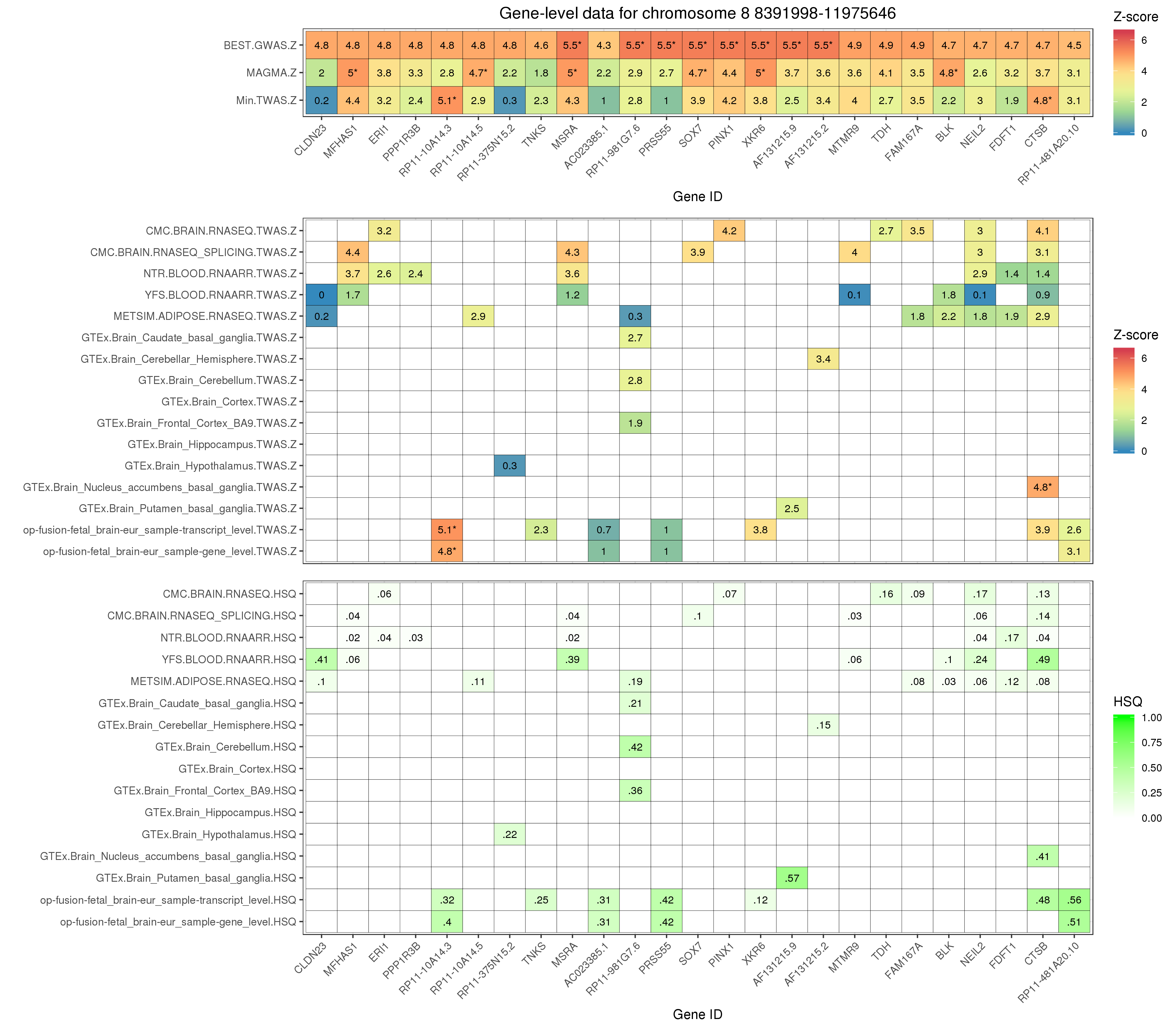

Supplementary Figure 8. Comparison of gene-level Z scores derived using MAGMA and TWAS, and SNP-level Z score in the corresponding GWAS, containing either a significant TWAS or MAGMA association. The absolute TWAS Z score is used here. Genes within 250kb of genes significant in the either the TWAS of MAGMA analysis are included. Empty cells indicate the gene was not available in the TWAS or MAGMA analysis. Asterisks indicate the Z score surpassed the corresponding significance threshold (TWAS = p < 4.25x10^-6^; MAGMA = Bonferroni p-value </= 0.05, GWAS = p <=5x10^-8^).

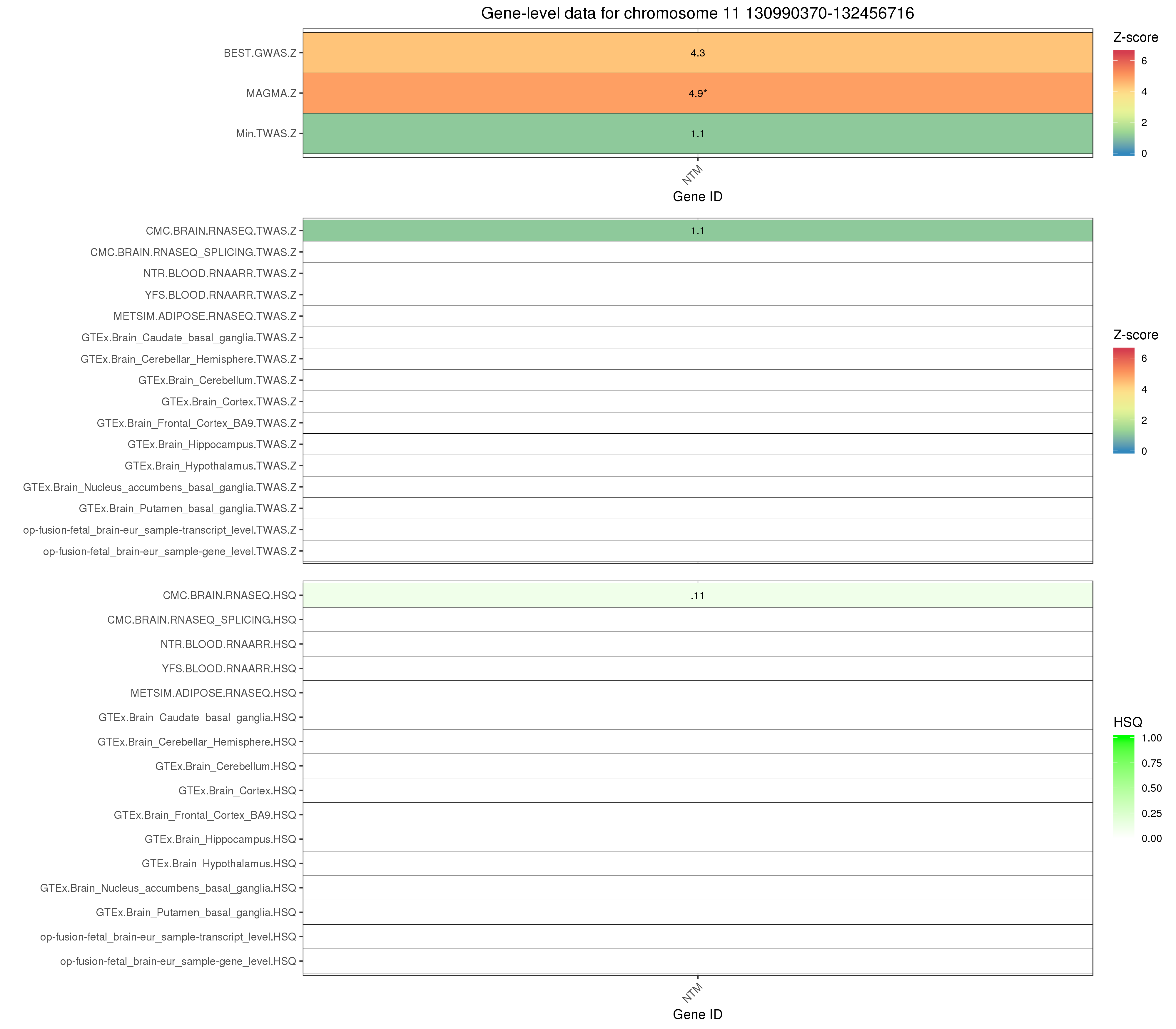

Supplementary Figure 9. Comparison of gene-level Z scores derived using MAGMA and TWAS, and SNP-level Z score in the corresponding GWAS, containing either a significant TWAS or MAGMA association. The absolute TWAS Z score is used here. Genes within 250kb of genes significant in the either the TWAS of MAGMA analysis are included. Empty cells indicate the gene was not available in the TWAS or MAGMA analysis. Asterisks indicate the Z score surpassed the corresponding significance threshold (TWAS = p < 4.25x10^-6^; MAGMA = Bonferroni p-value </= 0.05, GWAS = p <=5x10^-8^).

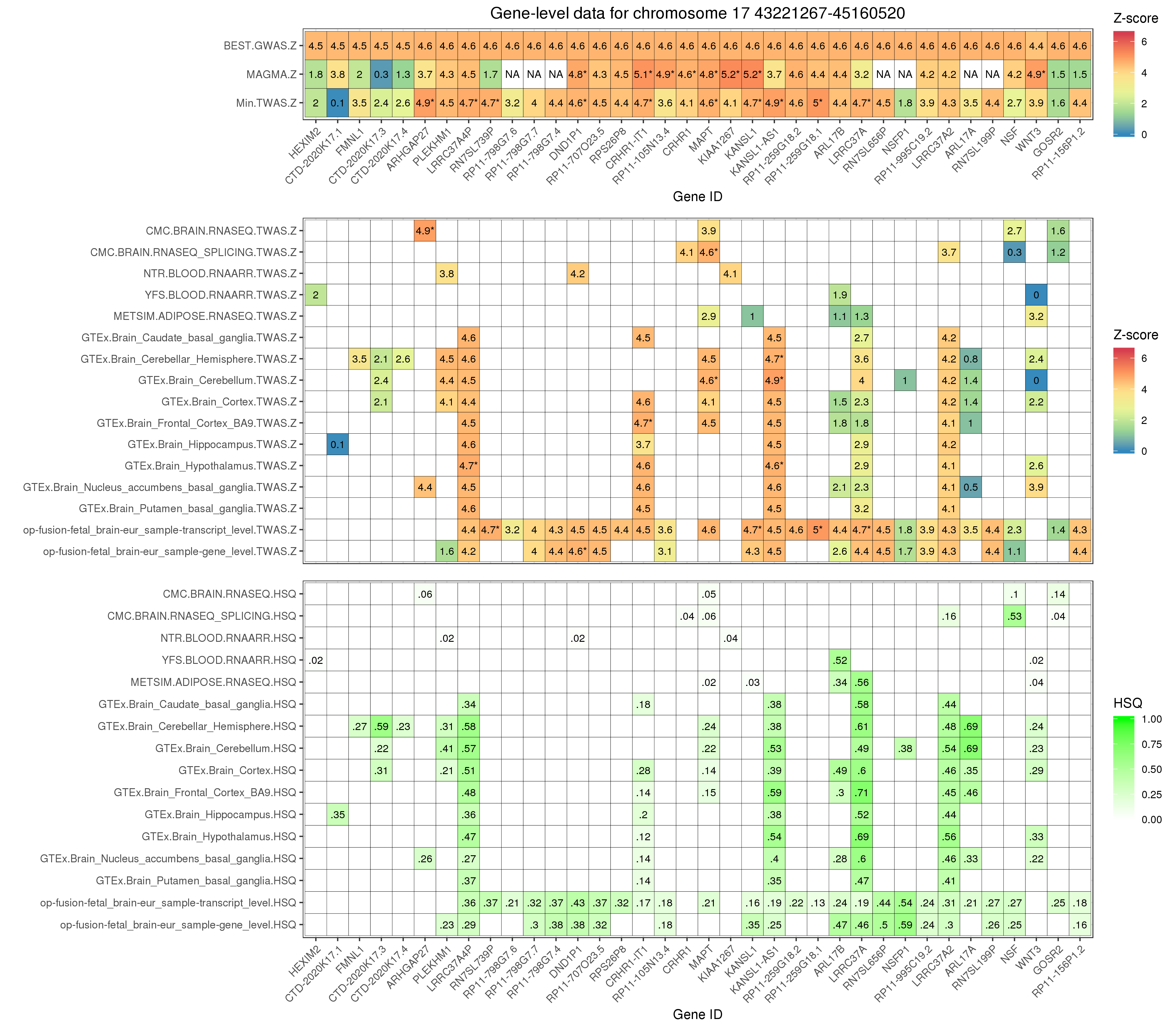

Supplementary Figure 10. Comparison of gene-level Z scores derived using MAGMA and TWAS, and SNP-level Z score in the corresponding GWAS, containing either a significant TWAS or MAGMA association. The absolute TWAS Z score is used here. Genes within 250kb of genes significant in the either the TWAS of MAGMA analysis are included. Empty cells indicate the gene was not available in the TWAS or MAGMA analysis. Asterisks indicate the Z score surpassed the corresponding significance threshold (TWAS = p < 4.25x10^-6^; MAGMA = Bonferroni p-value </= 0.05, GWAS = p <=5x10^-8^).

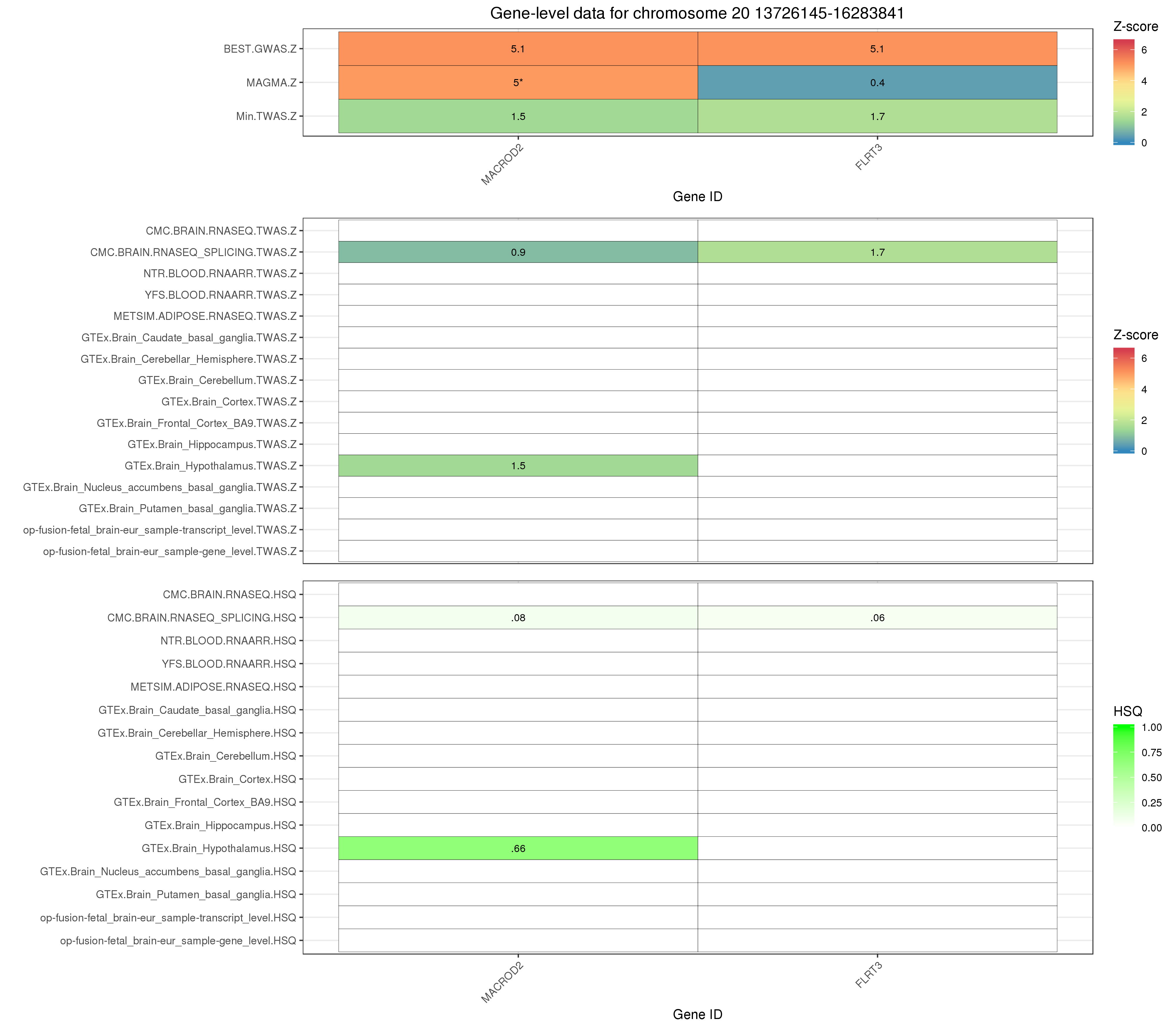

Supplementary Figure 11. Comparison of gene-level Z scores derived using MAGMA and TWAS, and SNP-level Z score in the corresponding GWAS, containing either a significant TWAS or MAGMA association. The absolute TWAS Z score is used here. Genes within 250kb of genes significant in the either the TWAS of MAGMA analysis are included. Empty cells indicate the gene was not available in the TWAS or MAGMA analysis. Asterisks indicate the Z score surpassed the corresponding significance threshold (TWAS = p < 4.25x10^-6^; MAGMA = Bonferroni p-value </= 0.05, GWAS = p <=5x10^-8^).

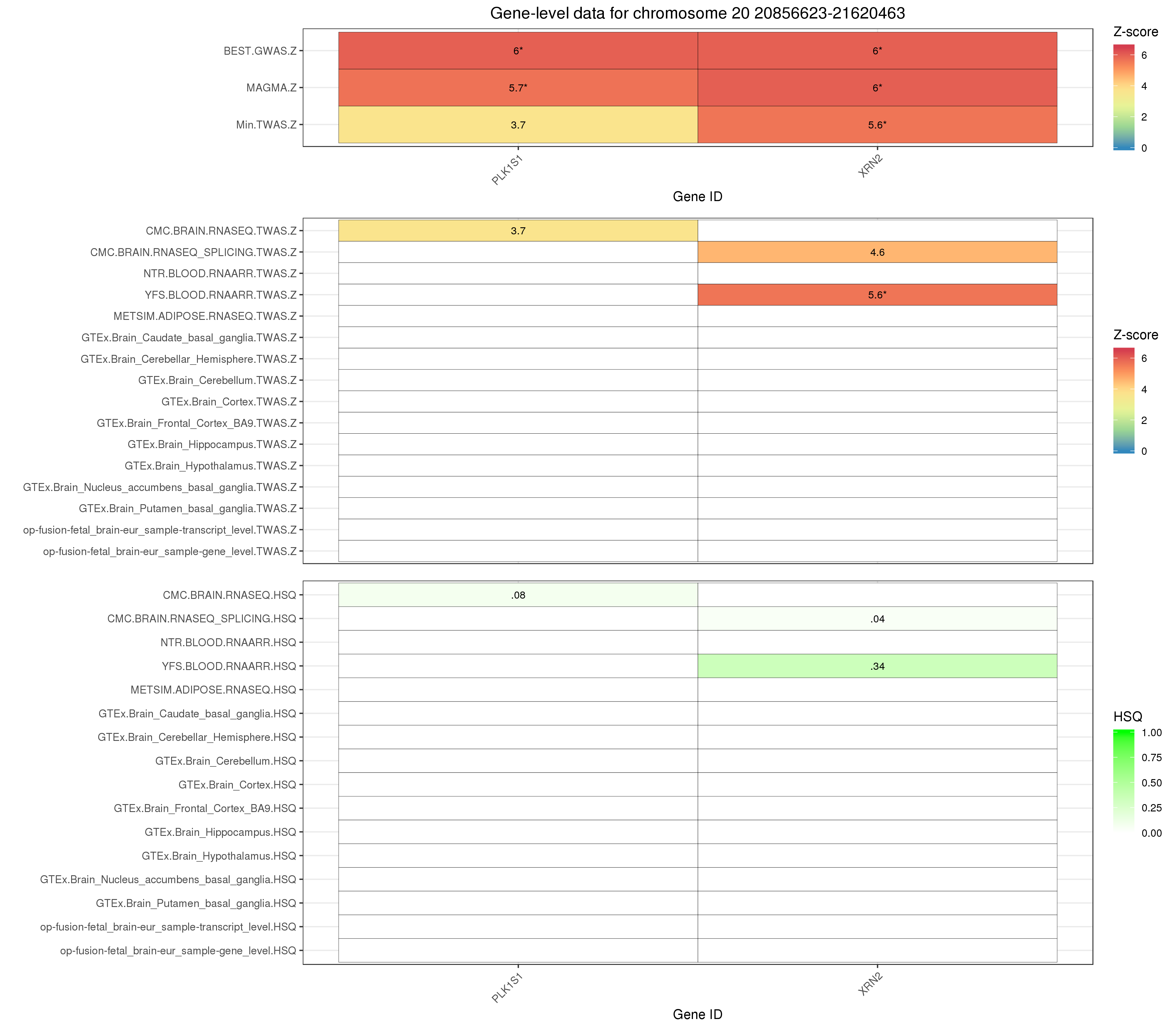

Supplementary Figure 12. Comparison of gene-level Z scores derived using MAGMA and TWAS, and SNP-level Z score in the corresponding GWAS, containing either a significant TWAS or MAGMA association. The absolute TWAS Z score is used here. Genes within 250kb of genes significant in the either the TWAS of MAGMA analysis are included. Empty cells indicate the gene was not available in the TWAS or MAGMA analysis. Asterisks indicate the Z score surpassed the corresponding significance threshold (TWAS = p < 4.25x10^-6^; MAGMA = Bonferroni p-value </= 0.05, GWAS = p <=5x10^-8^).
